# Peak performance is repeatable and captures large individual differences in ruby-throated hummingbirds

**DOI:** 10.64898/2026.05.29.728840

**Authors:** Émilie Gagnon, Juan Camilo Ríos-Orjuela, Lacey Pilon, Percy Hentschel, Alexa Dansereau, Paolo S. Segre, Roslyn Dakin

## Abstract

Locomotor performance often determines the outcome of interactions with competitors, predators, and prey. In flying animals, the asymptotic load-lifting assay measures maximal muscle power output in vertical flight. Previous studies of small birds have shown that load-lifting performance is linked to flight maneuverability and the outcome of competitive species interactions. Here, we quantify sources of performance variation within a species, namely repeatability, and determine the number of trials that accurately capture individual differences. We conducted 124 load-lifting trials on 13 wild-caught male ruby-throated hummingbirds (*Archilochus colubris*), testing each individual repeatedly over a 3-day period. We report large individual differences in peak performance, with 70% of the total variation in lifted mass attributed to differences among males. Notably, these differences in muscle performance are independent of body mass and size. An additional 23% of the variation in lifted mass was due to short-term fluctuations, wherein a given male’s performance varied across trials and days. We find no systematic effects of experience or time on load-lifting performance. Using simulation, we test the effect of different sampling protocols for measuring individual performance, and show that single-trial protocols yield the highest repeatabilities, but are less suitable for capturing the true underlying differences among individuals. We discuss recommendations for future studies that aim to measure maximum performance. Overall, our results show that the asymptotic load-lifting assay reveals large individual differences and can closely reflect individuals’ true maximum capacity.

**Summary statement:** Male ruby-throated hummingbirds exhibit large, consistent individual differences in load-lifting performance that are not explained by differences in body mass nor skeletal size.

## Introduction

Motor performance, defined as one’s ability to use its body to complete an ecologically relevant task (Irschick and Higham, 2015), can determine fitness through success or failure in predation, competition, escape, and dispersal (Domenici et al., 2007; Husak and Fox, 2008; Wilcox and Clark, 2022). According to the morphology-performance-fitness paradigm, an animal’s morphology and physiology will limit its performance. When competing, an animal will use its performance ability – to the extent that its physiology allows – to achieve survival and reproduction (Arnold, 1983; Irschick, 2003; Irschick and Garland, 2001). When fitness advantages depend on motor performance, heritable differences between individuals in the traits that underly that performance are expected to be under selection (Garland and Carter, 1994).

Consequently, a trait that differs consistently among individuals may also generate differences in survival and reproduction (Herde and Eccard, 2013; Réale et al., 2010).

When a performance trait influences competitive outcomes, the extent to which competitors differ is expected to influence the costs and benefits of assessment and escalation. For example, traits with larger among-individual differences tend to generate more consistent competitive asymmetries, where escalations are less frequent (Parker, 1974). By contrast, traits with small individual differences may generate longer conflicts, with greater potential for escalation of intensity (Bridge et al., 2000; Parker, 1974), and may allow for competitive outcomes to be determined by other social factors (Hardy and Briffa, 2013). When testing the relationship between locomotor performance and competitive outcomes, a key assumption is that performance assays accurately capture consistent differences among individuals (Garland et al., 1990). However, the extent of individual differences in biomechanical performance assays is not often reported. Furthermore, it remains challenging to determine whether a performance assay accurately captures underlying individual abilities (Careau and Garland, 2012).

One of the most widely accepted ways to assess the extent of individual differences is to measure performance from each individual repeatedly through time, and use a variance partitioning analysis (Bell et al., 2009; McGraw and Wong, 1996). Using a mixed-effects regression model with performance measure as the response variable, individual repeatability is calculated as the proportion of total variance attributed to differences among individuals, ranging from 0 to 1. Repeatability values closer to 1 indicate larger (or stronger) among-individual differences (McGraw & Wong, 1996; Nakagawa & Schielzeth, 2010). Repeated measures also allow testing the effects of time and experience. Performance may improve over short-term time through training or habituation effects (Davison, 1997; Owerkowicz and Baudinette, 2008); performance may also worsen, as a result of fatigue (Barry and Enoka, 2007; Fitts, 1994). These effects vary depending on the experimental design and animal model (Garland et al., 1987; Gleeson, 1979; Zug, 1985).

In this study, our aim was to determine the nature and extent of maximal flight performance differences among wild-caught male ruby-throated hummingbirds (*Archilochus colubris*). Hummingbirds are small nectar-feeding birds (family Trochilidae) that hover to feed (Powers and Nagy, 1988; Suarez, 1992). They can require hundreds of nectar meals per day (Wolf and Hainsworth, 1977), and as a result, many hummingbird species compete fiercely over access to flowers, with frequent chases and aggression occurring in flight (Feinsinger and Chaplin, 1975; Sargent et al., 2021; Sholtis et al., 2015). Flight performance is also essential for escaping threats (Lima, 1993; Miller and Gass, 1985) and, in many hummingbird species, for courting potential mates (Clark et al., 2011; Hogan and Stoddard, 2018; Wilcox and Clark, 2025). Given this background, flight muscle performance is expected to be an important determinant of survival and reproductive success in hummingbirds (Altshuler, 2006; Altshuler et al., 2004a; Altshuler et al., 2004b; Sargent et al., 2021; Segre et al., 2015; Sholtis et al., 2015; Wilcox and Clark, 2025).

The asymptotic load-lifting assay is widely used to measure maximal burst muscle capacity during vertical flight (Chai and Millard, 1997; Chai et al., 1997; Marden, 1987), and is readily used in hummingbirds as well as insects, bats, and other bird species (Araya-Salas et al., 2018; Buchwald and Dudley, 2010; Davis and Cockrum, 1965; Fernández et al., 2017; Kou et al., 2022). This assay is designed such that an animal in vertical flight must lift increasing mass until it reaches its maximal capacity (Chai et al., 1997; Fig. 1A). The mass can either be lifted via a thread with evenly spaced load units (Fig. 1A), or mass can be sequentially added, though the latter method is expected to underestimate maximal performance (Buchwald and Dudley, 2010). In hummingbirds, peak performance on this assay has been linked to species and individual differences in ecology and behaviour (Altshuler, 2006; Altshuler et al., 2004a; Altshuler et al., 2004b; Dakin et al., 2018; Segre et al., 2015). Previous studies using this assay typically report only one measurement per individual (see Table S1 for details). Only two previous studies have reported among-individual differences in load-lifting performance, based on repeated measures (Araya-Salas et al., 2018; Najar et al., 2026). Those two studies, done on two other hummingbird species, report substantial differences among males (repeatabilities 0.30 – 0.90) that were tested shortly after capture, with the repeated measures occurring within minutes of the first test (Najar et al., 2026) or upon second capture at varying intervals (Araya-Salas et al., 2018).

**Fig. 1.**
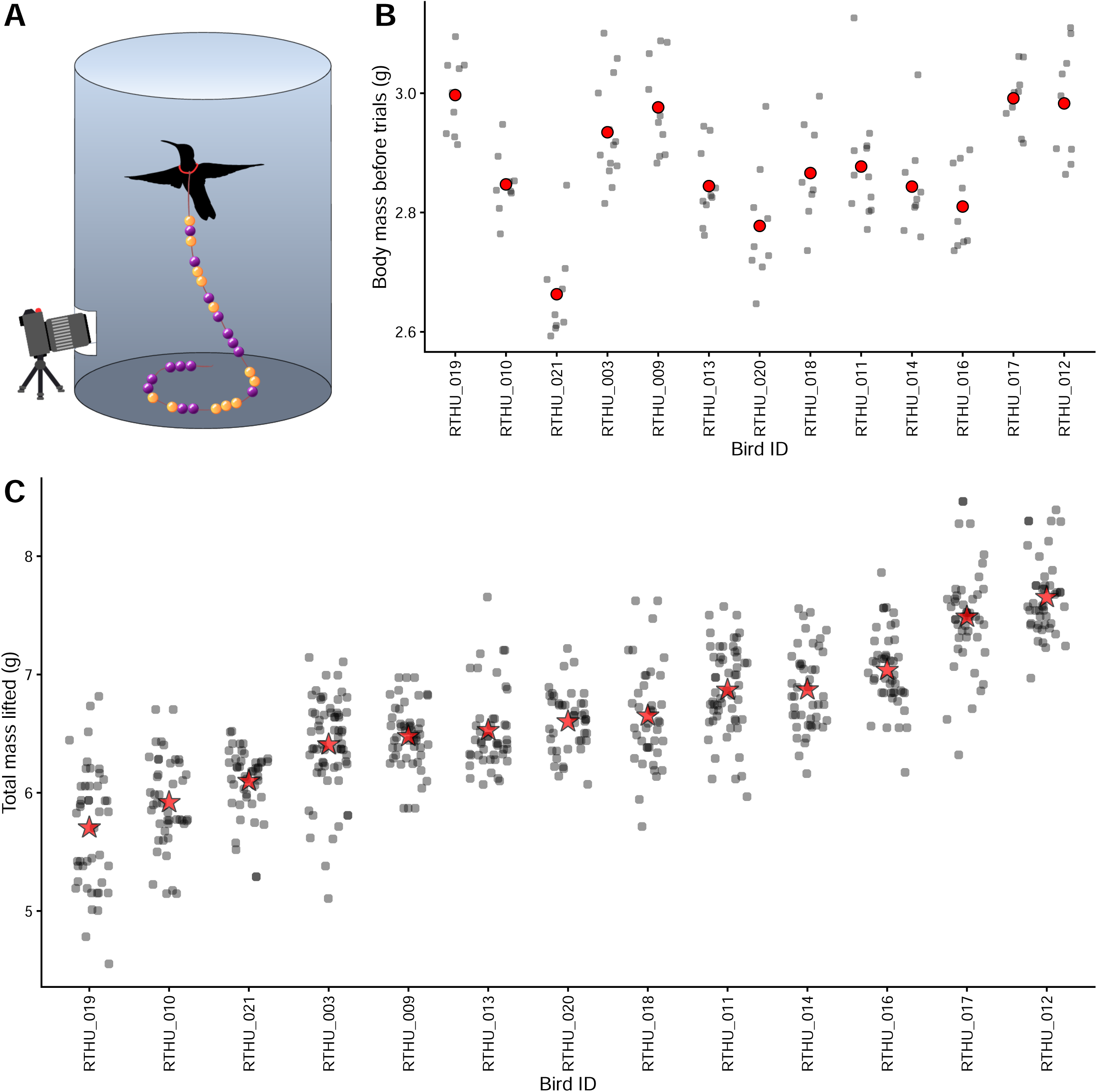
The load-lifting assay measures burst muscle performance in ruby-throated hummingbirds. (A) Diagram of the asymptotic load-lifting assay for hummingbirds. A bird is released from the bottom of a cylinder that is open at the top. As the bird flies vertically, it lifts more mass. The bead units each have a unique color code that facilitates scoring. Note that features in (A) are not to scale. (B) Body mass of 13 male ruby-throated hummingbirds before each load-lifting trial. Each black point represents a body mass measurement, with individual averages in red. (C) Load-lifting values (*Lift_total_*) are represented as grey points and each bird’s average is indicated by a star. In (B) and (C), the birds are ordered on the x axis by their average load-lifting performance.

Here, our aim was to determine the sources of variation in maximum performance within a species. We captured 13 adult male ruby-throated hummingbirds on their breeding grounds after migration and tested each male in the load-lifting assay repeatedly and systematically over a 3-day period, under the same conditions. Given that we are considering within-species differences, we use the peak mass lifted (per lift) as the focus of our analyses (Altshuler, 2006; Segre et al., 2015). We also examine a relative measure, the number of body mass units lifted, as a secondary analysis presented in the supplement. We show strong agreement for peak load-lifting measures within the same trial. We then use variance partitioning for our primary aim of quantifying sources of variation in peak performance, and evaluating whether load lifting is determined by a male’s body mass and size. We further evaluate the extent to which individual performance fluctuates across and within days. Finally, we develop a simulation framework to test the effect of different sampling protocols for determining individual differences in underlying maximum ability. Based on our findings, we provide general guidelines for accurate animal peak performance assays.

## Materials and Methods

### Animal capture and housing

We captured 13 adult male ruby-throated hummingbirds (*Archilochus colubris*) in Eastern Ontario in May and June 2025. Upon capture, birds were housed indoors in individual cages (95 cm x 45 cm x 45 cm) with *ad libitum* artificial nectar solution (Nektar-plus, Nekton). The holding facility was held at 22°C and 30-60% relative humidity, with a daily light cycle of 13h light:11h dark. All procedures were approved by the Carleton University Animal Care Committee.

### Load-lifting apparatus

The aim of the asymptotic load-lifting assay is to elicit maximum effort in vertical flight. As hummingbirds are highly aerial and visually guided, their escape response is to flee from locations that are enclosed, on the ground, or darkened, and escape towards areas that are open, raised, and bright. The load-lifting apparatus was based on previous studies (Chai et al., 1997; Mahalingam and Welch, 2013) and exploits hummingbirds’ strong response to the three features listed above, so that the assay consistently elicits the intended response of vertical escape flight. The apparatus for the assay was a smooth paper cylinder (45 cm wide x 70 cm high) with an open top (Fig. 1A). We chose the dimensions of the cylinder to minimize ground and boundary effects, with height > 50-60 cm and width greater than four times the animal’s wingspan, following Rayner and Thomas (1991). We used a small door on the side of the cylinder side to position birds inside. We used an opening at the bottom of the cylinder to film the interior using a GoPro HERO8 camera, filming at 120 fps (1080p and automatic shutter speed; Fig. 1A). We installed a second camera (Wyze Cam v3, filming at 1080p and 20 fps) above the cylinder to monitor the assay in real time.

To construct the load to be lifted by the hummingbirds, we used a nylon thread (96 cm long) with 32 load units, spaced 2.0 cm apart (Fig. 1A). Each load unit was composed of 5-6 beads, with a mass of 0.150 or 0.229 g per unit including the mass of the thread. Bead unit mass and spacing were chosen based on literature recommendations (Rayner and Thomas, 1991) and previous load-lifting data conducted on ruby-throated hummingbirds (Chai et al., 1997; Mahalingam and Welch, 2013). To facilitate scoring the total amount lifted, we used distinct color combinations in each load unit. We used a small elastic loop (0.088 g) at one end of the thread to mount the load around the bird’s neck (Chai et al., 1997). In total, the thread (including all bead units and the elastic) weighed 5.898 g.

We assume that the load-lifting assay elicits maximum effort, because it combines a natural escape response with an asymptotically increasing load. Three further observations support our expectation that the hummingbirds were motivated to perform with maximum or near maximum effort in this assay. First, during the asymptotic load-lifting trials, the ruby-throated hummingbirds gaped, which is a sign of extreme ventilatory demand in birds (Brackenbury, 1984). Second, during preliminary testing in this study, we checked whether hummingbirds would perform better on this assay after they had been permitted to freely escape the cylinder without the loaded thread. We found no difference in asymptotic load-lifting performance before or after a free escape, suggesting that motivation to escape in this assay is not readily altered (see supplement for details). Third, in a related hummingbird species, *Calypte anna*, the ceiling of aerodynamic force produced in the highest-performance arcing turns is consistent with the peak aerodynamic force generated vertically in the load-lifting assay (Segre et al., 2025).

### Load-lifting trials

We aimed to test birds under equivalent laboratory conditions, as close to their wild-caught state as possible. To achieve this, we ran load-lifting trials on their 2^nd^, 3^rd^, and 4^th^ morning after capture, allowing the first full day after their capture day for acclimation and to ensure that all birds fed *ad libitum* for a full day prior to testing. Each bird was tested in three separate bouts on a given test day (hereafter, ‘trials’), with a rest period in between each trial in which the bird was returned to its home cage for 30 minutes. Trials took place between 8:00 h and 13:00 h. We randomized the time start of each bird’s 1^st^ trial of the day.

To initiate a trial, we placed the load around the bird’s neck and positioned the bird in the middle of the floor of the cylinder (Movie 1). We arranged the thread around the bird in a spiral shape to avoid tangling. The bird was then released and the cylinder door quickly closed. After a bird completed a series of lifts, we allowed the bird to rest on the floor of the cylinder for at least 30 seconds (Chai et al., 1997), after which we repositioned the bird and thread to the starting position. A trial was ended when a bird performed at least five vertical lifts, as monitored from the top camera. In a few cases, we conducted an additional trial and/or test day when a bird could not perform the assay (*n* = 8 extra trials conducted). To score the trials, we reviewed the GoPro video frame-by-frame. We only scored lifts in which the loaded thread was lifted vertically and did not touch the cylinder or the experimenter’s hands. For each scored lift, we recorded the final load unit raised from the floor. Then, from each trial, we retained the five best lifts as ranked by the number of load units lifted (see supplement for details) and computed the mass of the beaded thread lifted in each of the five best lifts per trial (i.e., mass of the elastic loop, thread, and bead units lifted above the floor).

### Body mass and skeletal size measurements

Before and after each load-lifting trial, we measured the bird’s body mass to the nearest 0.001 g (Ohaus Scout SPX123; Fig. 1B). Prior to conducting the load-lifting trials with a given male, we took three measures of skeletal size: keel length, exposed culmen length (hereafter, ‘bill length’) and tarsus length (Freeman and Jackson, 1990; Peig and Green, 2010; Senar and Pascual, 1997). We measured tarsus length to the nearest 0.1 mm using a digital caliper (Simhevn electronic caliper). For bill and keel length, we used macrophotography with a standardized scale in each image (Canon EOS M50, 162 mm macro lens). We used the first principal component scores from a PCA of keel, bill, and tarsus measurements as an overall measure of a bird’s skeletal body size (see supplement and Tables S2-S3 for details).

### Analysis of load-lifting performance

All data analyses were performed using R v4.5.1 (R Developmental Core Team 2025). We focused our main analyses on total mass lifted (*Lift_total_*) as a measure of overall force production. We calculated *Lift_total_* for each lift as the mass of beaded thread lifted, plus the bird’s trial-average body mass (i.e., average of its pre– and post-trial body mass).

As a check on measurement agreement, we ran a concordance correlation for the two best *Lift_total_* values from each bird’s first daily trial (*n* = 39 trials). We expect a fresh bird in its first trial of the day to be as close as we can obtain to a fixed subject. We used Lin’s concordance correlation, which yields two estimates. First, the bias correction factor C_b_ ranges from 0-1, wherein 1 indicates a complete absence of systematic bias (or scale shift) between two measures. Second, the concordance correlation coefficient, CCC, is the product of C_b_ (accuracy) and Pearson’s r (precision). A value of Lin’s CCC near 1 would indicate that the two best *Lift_total_* values from each trial are in near-perfect agreement.

Our primary aim was to investigate sources of variation in peak lift performance. To do this, we fit generalized linear mixed-effect regression models (GLMMs) with a Gaussian error distribution using the R package *lme4* v1.1-37 (Bates et al., 2015). We modelled lift performance as a function of five fixed effect predictors that we expected to influence performance, as well as three random effects to account for the structure of our repeated measures of birds, days, and trials:

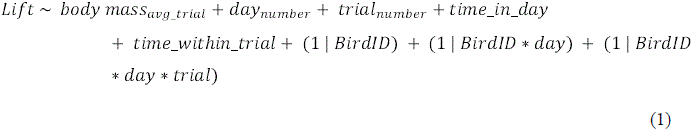

The random effect of bird ID accounts for individual differences in performance (repeatability), whereas the effect of bird ID * day accounts for consistent variation in performance by a given bird among days, and bird ID * day * trial accounts for consistent variation among a bird’s trials. The model above (Eqn 1) includes a fixed effect predictor for a bird’s average trial body mass (‘body mass_avg_trial_’), as well as two fixed effect predictors for a bird’s prior experience in the assay: ‘day_number_’, accounting for changes across days, and ‘trial_number_’, accounting for changes across consecutive trials within a day. The predictor ‘time_in_day’ captures potential changes with time of day in minutes (Kusumoto et al., 2021), whereas the predictor ‘time_within_trial’ captures potential changes over time elapsed within a given trial (in seconds). We included these temporal effects to account for possible changes in performance driven by training and/or fatigue. We standardized all fixed effect variables, with mean 0 and standard deviation of 1. We used the R package *DHARMa* v0.4.7 (Hartig, 2024) to verify that model residuals were normally distributed and homoscedastic.

To characterize the major sources of variation in lift performance, we calculated the proportion of total variance in lifting performance attributed to each random effect using the R package *rptR* v0.9.23 (Nakagawa and Schielzeth, 2010; Stoffel et al., 2017). The proportion of variance attributed to bird ID represents repeatability, or the extent of among-individual differences accounting for other effects in the fitted model (McGraw and Wong, 1996; Nakagawa and Schielzeth, 2010). The proportion of variance attributed to bird ID * day indicates the extent to which performance is explained by day-to-day fluctuations for a given bird. Likewise, the proportion of variance attributed to bird ID * day * trial captures trial-to-trial fluctuations. We used parametric bootstrapping (1,000 iterations) to obtain 95% confidence intervals (CIs) on these estimates.

To focus on a bird’s maximal performance, we present results using a dataset that included only the two best lifts per trial, but we also ran two alternative models. We repeated analyses using all datapoints (five best lifts per trial) to determine whether a larger number of sampled lifts per trial alters the repeatability estimate and whether this increased sample size could reveal fixed effects that would not be significant in the model with two best lifts only. We also examined a model of mass-specific lifting performance (*Lift_relative_*; two best lifts per trial), which is calculated as the total mass lifted divided by the bird’s trial-average body mass. Note that for the *Lift_relative_* model, we removed body mass as a predictor.

### Effect of sampling on the measurement of individual differences

To determine the number of load-lifting trials required to accurately measure individuals’ best *Lift_total_* performance, we sub-sampled our data to generate different sampling protocol scenarios wherein the assay was performed over fewer days and trials (Fig. S1). For each scenario, we randomly sampled the required number of days and trials from each bird, and retained the two best lifts per trial. To simplify comparison across all sub-sampling protocols, we used a simplified model of lift performance shown in Eqn 2 below to calculate repeatability. We repeated this procedure 500 times to generate a distribution of repeatability estimates for each scenario.

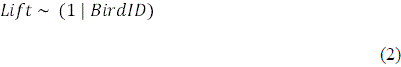

To test whether the highest repeatability sampling protocol also gives the most accurate measure of individual performance, we used a simulation, generating data for *n* = 13 *in silico* individuals wherein their true maximal capacity is known. We defined an individual’s true maximal capacity as the sum of its underlying individual mean and 1.5 times the other sources of variation (daily variation, trial variation, and residual errors), giving an estimate of “best possible performance” for that individual. Note that we used 1.5 as the constant here because the resulting variance component magnitudes were consistent with those observed in our empirical hummingbird data. We assume that individuals differ in muscle capacity (underlying individual means), and that performance on any one lift is generated by these underlying individual mean differences, plus day-to-day fluctuations in performance, trial-to-trial fluctuations in performance, and measurement error. We used means and variances derived from our real (empirical) hummingbird data to generate lift events with these sources of variation (see Fig. S2 for details). For each *in silico* individual, we generated a sample size of five lifts per trial, three trials per day, and three days. After generating the simulated lifts, we performed the same sub-sampling protocol described above, randomly selecting 1-3 days and trials. From each sub-sampling protocol (*n* = 500 sampling iterations per protocol), we calculated (i) repeatability (Eqn 2), and (ii) the correlation between the individuals’ true maximal capacity and their estimated maximal performance, with their estimated max defined as the average of their two maximal lifts in each sampled trial.

## Results

We recorded 614 best lifts for 13 birds across 124 trials (five lifts per trial, range of trial duration = 3-13 minutes; Fig. 1C). In four trials, the bird performed fewer than five lifts. The average peak performance was 2.4x body mass (range 1.5-2.9x body mass). This is consistent with mass-specific lift performance for a congener hummingbird species, the black-chinned hummingbird *Archilochus alexandri*, tested within an hour of capture (Wilcox and Clark, 2025).

### Measurement agreement

We find very strong agreement, with a concordance CCC of 0.93 (95% CI 0.86-0.96) and a bias correction factor C_b_ > 0.99 for peak load-lifting samples within a trial.

### Load-lifting performance

Repeatability of *Lift_total_*was high, at 0.70 for the ‘two best lifts per trial’ protocol, indicating that individual differences that are consistent across days explain 70% of the total variation in peak performance (Fig. 2A and Table 1; *n* = 248 lifts in 124 trials). An additional 12% of the variation in *Lift_total_* is attributed to fluctuations for a given bird across days, and an additional 11% is attributed to fluctuations for a given bird across trials (Fig. 2A and Table 1; all components explain statistically significant variation in performance). These latter two estimates indicate that individual males have periods in which they perform consistently better (or worse) relative to their average performance (Fig. 2B-D).

**Fig 2.**
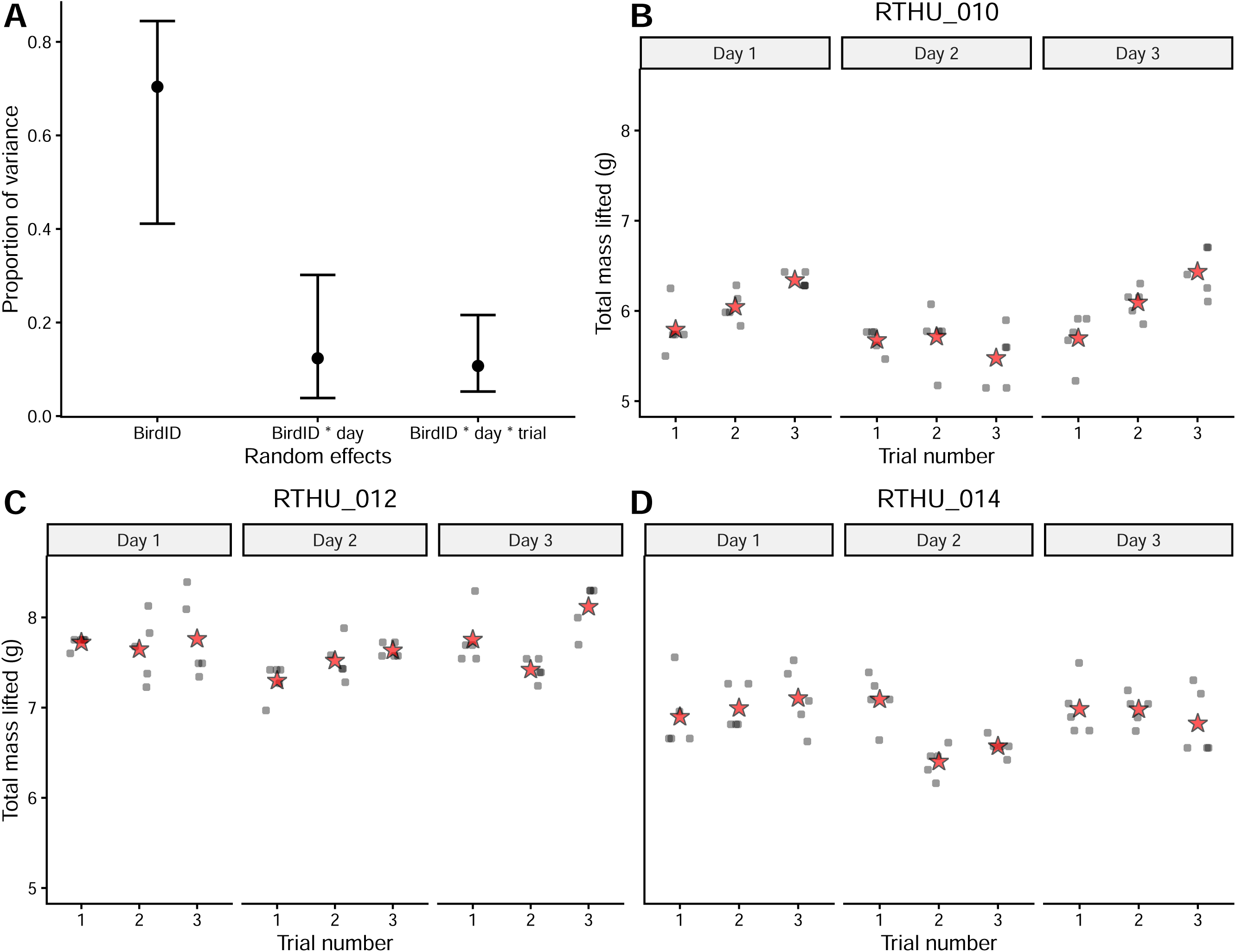
Individual males differ consistently in peak performance (*Lift_total_*), with additional variation across and within days. (A) Proportion of variance in *Lift_total_* explained by differences among individuals (BirdID), consistent day-to-day differences for a given individual (BirdID * day), and consistent trial-to-trial differences for a given individual (BirdID * day * trial). Error bars show 95% confidence intervals from bootstrapping (*n* = 1,000 iterations). (B-D) Examples illustrating how *Lift_total_* varies across and within days for three hummingbirds: RTHU_010, RTHU_012, and RTHU_014. Each gray point represents a lift, and each star represents the average per trial.

**Table 1.**
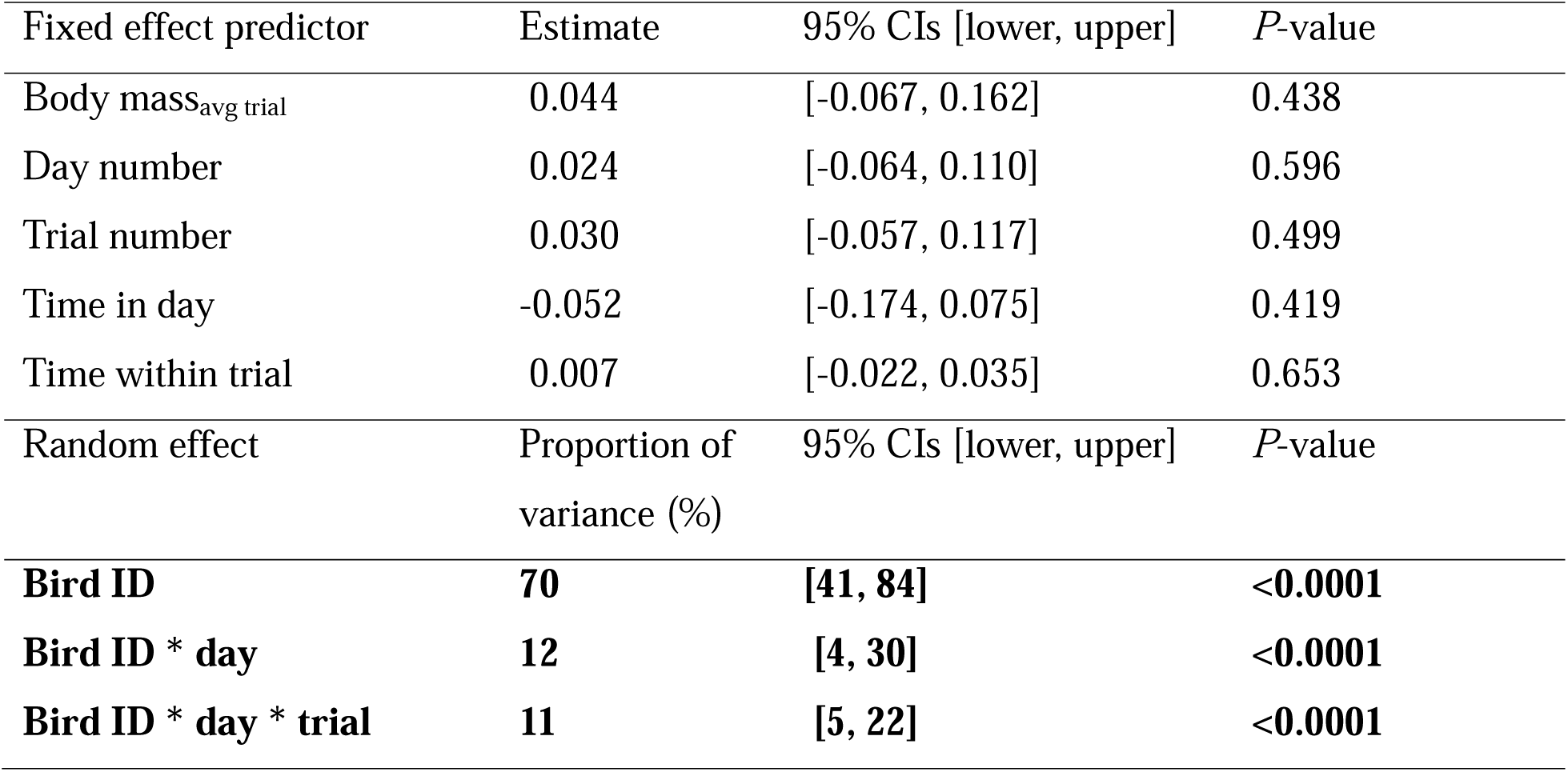
Analysis of lifting performance using the two best lifts per trial. The dependent variable is *Lift_total_*. All fixed effect predictors were centred and standardized, so that the coefficient estimates here are comparable. Variables are bolded when *P*-values are below 0.05.

| Fixed effect predictor | Estimate | 95% CIs [lower, upper] | $P$-value |
| --- | --- | --- | --- |
| Body mass<sub>avg trial</sub> | 0.044 | [-0.067, 0.162] | 0.438 |
| Day number | 0.024 | [-0.064, 0.110] | 0.596 |
| Trial number | 0.030 | [-0.057, 0.117] | 0.499 |
| Time in day | -0.052 | [-0.174, 0.075] | 0.419 |
| Time within trial | 0.007 | [-0.022, 0.035] | 0.653 |
| Random effect | Proportion of variance (%) | 95% CIs [lower, upper] | $P$-value |
| **Bird ID** | **70** | **[41, 84]** | **<0.0001** |
| **Bird ID \* day** | **12** | **[4, 30]** | **<0.0001** |
| **Bird ID \* day \* trial** | **11** | **[5, 22]** | **<0.0001** |

We did not detect significant effects of predictors related to a bird’s experience in the assay, time of day, or time elapsed within a trial on *Lift_total_* (Table 1; see also Fig. S3). We also did not find a significant effect of a bird’s body mass on his load-lifting performance (Fig. 1B, Fig. 3 and Table 1). To confirm this result using a different measure, we tested for an association between average individual *Lift_total_* and a bird’s skeletal body size (defined as the first principal component of keel, bill, and tarsus lengths). We found that skeletal body size is also not a significant predictor of an individual’s load-lifting performance (see Fig. S4 and supplement for details).

**Fig 3.**
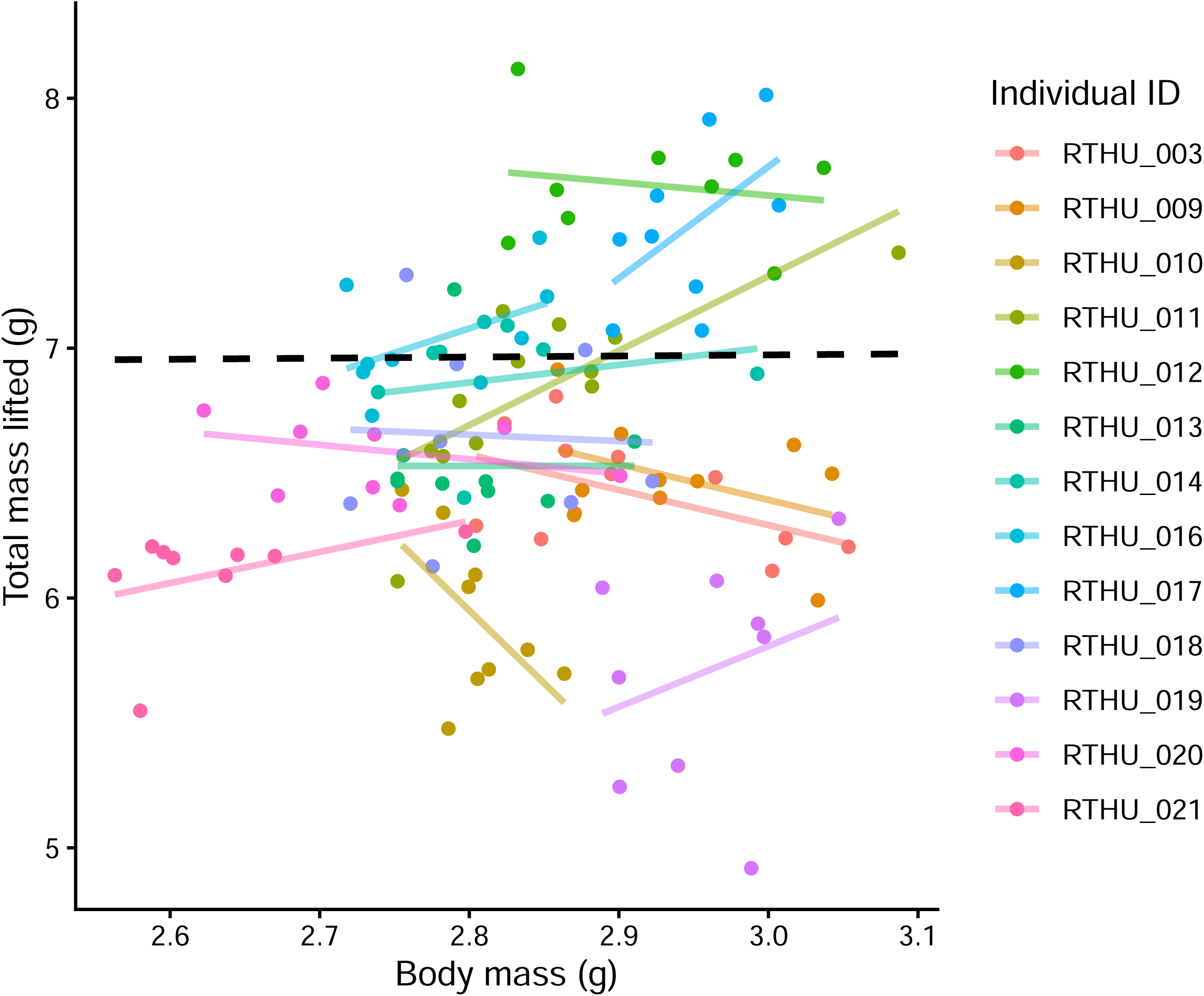
Individual variation in body mass does not predict differences in load-lifting performance among males. Each point represents the trial-average load-lifting performance of a male. Individual points and lines are colored by bird ID (*n* = 13). The black dashed line shows the ‘two best lifts per trial’ model prediction for the relationship between *Lift_total_*and bird body mass.

We reached the same general conclusions when analyzing performance using the ‘five best lifts’ protocol, or when analyzing *Lift_relative_* as the dependent variable. Briefly, the proportions of variance attributed to different sources in Fig. 2A are all within 1-5% of the estimates from the main model, and the effect sizes for the fixed effect predictors are also negligeable. These additional results are presented in the supplement (see Fig. S5 and Tables S4, S5 for details).

### Effect of subsampling protocol on the measurement of individual differences

Our estimates of repeatability (individual differences) in lift performance increased when the sampling protocol was limited to fewer trials or fewer days (Fig. 4A), with the highest repeatabilities found when sub-sampling only one trial per bird (average repeatability of 0.93).

**Fig. 4.**
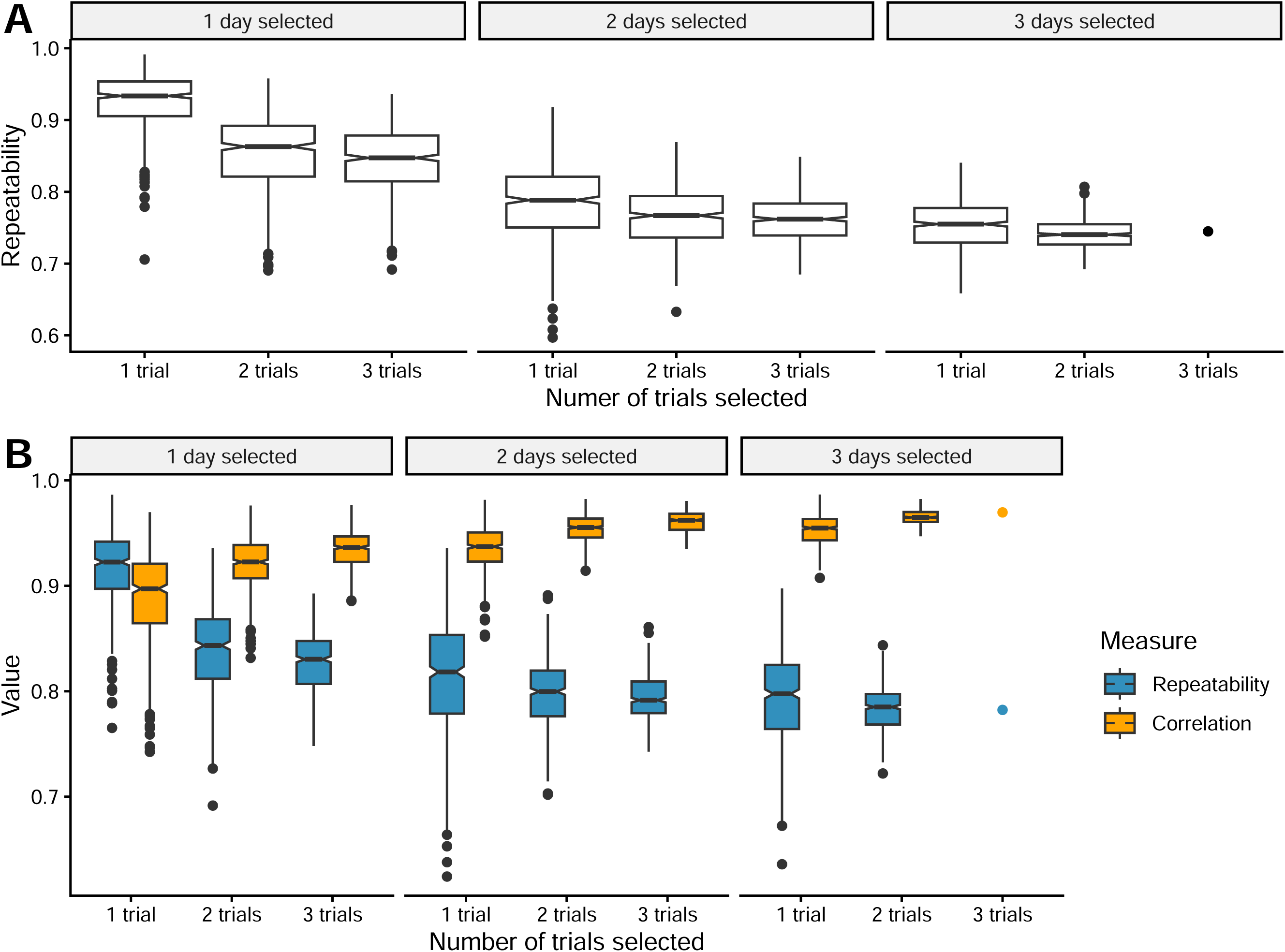
Effect of sampling protocol on repeatability and the accuracy of estimated performance maxima. (A) First, we sub-sampled our hummingbird load-lifting measurements to test the effect of limited sampling on repeatability. As shown in (A), repeatability is highest for protocols that are limited to fewer trials and fewer days. Second, we used a simulation in (B) to test how sampling protocols affect the accuracy of estimated individual performance maxima. Blue boxplots show repeatability estimates. Orange boxplots show estimates of the correlation between the individuals’ true maximal capacity and their sampled maximal performance. Each boxplot in A and B represents *n* = 500 iterations, with the exception of the protocol based on 3 days & 3 trials (*n* = 1). Boxplots are marked at the 25^th^, 50^th^ and 75^th^ percentiles, and the whiskers represent 1.5 x the interquartile range.

Our simulation demonstrates that although limited sampling protocols may yield higher repeatability, the estimates derived from them are a poorer indication of individuals’ true maximum capacity (Fig. 4B). As the number of trials included in the sampling protocol decreases, repeatability increases, but the correlation between the true underlying individual maximum and the sampled maximal performance decreases (from left to right in Fig. 4B).

## Discussion

Our study shows that male ruby-throated hummingbirds differ substantially in burst muscle capacity (Fig. 1C), across three days of repeated measures, with 70% of the variation in asymptotic load lifting attributed to among-individual differences (Table 1 and Fig. 2A). Previous studies of peak locomotor performance in other non-human animals report similar among-individual differences (Adolph and Pickering, 2008; Artacho et al., 2013; Hoskins et al., 2017; Jornod and Roche, 2015), including two studies of load lifting in other hummingbird species (Table S1; Araya-Salas et al., 2018; Najar et al., 2026). By testing individuals systematically across days here, we report that 23% of the variation in hummingbird peak performance was due to fluctuations in performance within a day (11%) and across days (12%) for a given individual (Table 1 and Fig. 2). In other words, 23% of performance variation was attributed to periods when a male would perform consistently better or worse than his overall average (e.g., Fig. 2B-D).

These short-term individual fluctuations are similar to the estimates reported in performance human studies, where performance varied within and between days at a ∼5-15% rate for each time scale (Banyard et al., 2017; Highton et al., 2012). These fluctuations may also be analogous to “hot/cold streaks” that have been reported in top-level human athletes (Bar-Eli et al., 2006; Gilovich et al., 1985; Reifman, 2011). To our knowledge, this is the first study investigating and thus reporting short-term individual fluctuations in performance in a non-human animal, beyond any systematic effects of time (e.g., fatigue or training effects; (Barry and Enoka, 2007; Davison, 1997; Fitts, 1994; Owerkowicz and Baudinette, 2008). Finally, we also found no consistent effects of time or a male’s amount of prior experience on performance across analyses (Table 1, Tables S4-5, and Fig. S3), indicating no clear time-of-day, fatigue, training, or habituation effects for the hummingbirds in this assay.

Individual differences in performance arise from genetic variation, environmental conditions, and the interaction of genes and environment throughout an animal’s lifetime (Houle, 1992; Mousseau and Roff, 1987; Roff and Mousseau, 1987). Our study focused on individual differences in male ruby-throated hummingbirds that had recently arrived from their long-distance pre-breeding migration of 2,000 – 3,000 km. Studies of body composition in other migratory bird taxa indicate that substantial flight muscle mass (∼25%) is consumed or lost during migration, followed by a recovery period when muscle mass increases (Battley et al., 2000; Bauchinger and Biebach, 2001; Groom et al., 2023; Karasov and Pinshow, 1998; Landys-Ciannelli et al., 2003; Pereira and Marini, 2015; Winker et al., 1992). In a study of migrant rufous hummingbirds (*Selasphorus rufus*), body mass increased by about 50% within 3 weeks at migration stopover sites, composed of an initial slow gain in non-lipid components (expected to be muscle), followed by a large and rapid increase in lipid stores (Carpenter et al. 1993). Although we know of no studies that directly measure muscle increases in ruby-throated hummingbirds, they too undergo large changes in body mass and body composition for migration (Zenzal and Moore, 2016).

In many migratory birds, males are selected for early arrival on breeding grounds, and individuals that arrive earlier are often in better condition (Marra et al., 1998; Møller, 1994; Neate-Clegg and Tingley, 2023; Rousseau et al., 2020). Given this background, a likely explanation for the large differences in muscle capacity among male ruby-throated hummingbirds here is the environmental conditions they experienced during their recent migration and arrival, and their ability to acquire resources for muscle gain post-migration. Thus, an important next step for improving our understanding of the sources of individual variation is to test whether body composition can explain individual differences in load-lifting flight (Dietz et al., 1999; Swanson and Merkord, 2013; Wilder et al., 2016). Given that fine-scale tracking of small birds (< 5g) is now also possible (Rhodes et al., 2025), future studies may also use these techniques to test how migration influences performance on the breeding grounds, with potential impacts on competition for territory and mates.

### Measuring peak performance

Our sub-sampling analysis showed higher repeatabilities for protocols with fewer trial samples (Fig. 4A). Our simulation results show how this is driven by consistent trial-to-trial fluctuations in the hummingbirds’ performance (Fig. 4B and Fig. 2B-D). Protocols with fewer trials will also decrease the accuracy of the sampled individual maximum (Fig. 4B). As the peak performance of a given male will vary across trials and days, sampling an individual more times leads to a higher probably of sampling near its true maximum (Gaines and Denny, 1993). Taken together, we advise caution when interpreting repeatability; high repeatability does not guarantee accurate measurements of individual capacities (Careau and Garland, 2012). If an assay generates short-term fluctuations or “streaks” in peak performance, we recommend integrating measurements across multiple time periods for a more accurate picture of true underlying individual capacities.

From our simulation, we also emphasize that the correlation between the individuals’ true maximal capacity and their estimated maximal performance remains high, even when sampling one trial per individual (mean correlation of 0.89; Fig. 4B). Such a high correlation suggests that the peak load-lifting assay can indeed capture values closer to individuals’ true maximal capacity, compared to non-peak or average locomotor performance (Banyard et al., 2017; Careau and Garland, 2012). Indeed, performance assays can either report peak performance or a behavioural trait, where the former quantifies what an animal *has* to do and the latter quantifies what an animal *chooses* to do (Careau et al., 2026). Below, we provide general guidelines to successfully conduct a peak performance assay that aims to measure true maximal capacities.

### Recommendations for study design

We recommend beginning with a literature review followed by preliminary testing. First, the experimental design of peak performance assays should be based on previous work done on the same (or a similar) assay. In this study, as mentioned in our methods, the asymptotic load-lifting assay was based on the pioneering work done by Chai and colleagues (1997), the aerodynamic recommendations formulated by Rayner and Thomas (1991), and by the previous studies of load-lifting data collected on the ruby-throated hummingbird specifically (Chai et al., 1997; Mahalingam and Welch, 2013). Second, a preliminary testing period should be used to ensure that the set up adequately triggers the intended response (Careau et al., 2026). Our preliminary tests allowed us to settle on opaque smooth walls, rather than perforated walls (see supplement). Third, if the variation among individuals is known in advance to be large, a single trial will capture these differences successfully (Fig. 4B). When possible, we recommend at least two well-separated trials per individual, because the sources of variation are typically not known in advance. Fourth, the assay must be ecologically relevant to the focal species and tailored to the study population (Careau and Garland, 2012; Irschick and Garland, 2001; Irschick et al., 2008). Hummingbirds are agile flyers that fight for resources on the wing, making load lifting an ecologically relevant assay as shown in cross-species hummingbird studies (Altshuler, 2006; Altshuler et al., 2004a; Altshuler et al., 2004b; Dakin et al., 2018).

### Ecological implications

In hummingbirds, flight maneuverability is hypothesized to determine survival and reproductive success (Altshuler, 2006; Altshuler et al., 2004a; Sargent et al., 2021; Sholtis et al., 2015; Wilcox and Clark, 2025). If peak flight performance determines dominance between hummingbird conspecifics (Altshuler, 2006; Altshuler et al., 2004a; Altshuler et al., 2004b; Sargent et al., 2021; Segre et al., 2015), the high repeatability that we report here for load lifting would suggest that competitive outcomes between males would be largely stable (Parker, 1974). We also report smaller, short-term fluctuations in individual performance, where males may experience hot or cold streaks. These fluctuations could contribute to short-term variation in competitive asymmetries, especially when two competitors have closely matched capacities. For instance, flight performance fluctuations could either give the advantage to an otherwise inferior individual, or further equalize fighting abilities of the two individuals and lead to conflict escalation (Bridge et al., 2000; Parker, 1974). Similarly, when two closely matched males compete over a resource, other traits might determine the winner, where for example, individuals with longer bills gain the advantage by stabbing their opponents in close contact (Araya-Salas et al., 2018; Rico-Guevara and Araya-Salas, 2015). For a more complete understanding of the drivers of dominance, future studies investigating dominance should analyze competitive asymmetries alongside traits that encompass both performance and traditional measures of morphology (Araya-Salas et al., 2018).

Thus far, whether hummingbird load-lifting performance correlates with courtship display performance remains uncertain. In the black-chinned hummingbirds (*Archilocus alexandri*), such a relationship was not found, though it is unclear whether the lack of correlation could be due to the courtship display comprising of lateral rather than vertical movement, or whether males are not performing at their maximal flying abilities during courtship (Wilcox and Clark, 2025). It would be interesting to test the relationship between load-lifting performance and courtship display performance in a species where male displays include more vertical flight components, such as the dive displays found in the Anna’s hummingbird (*Calypte anna;* Clark, 2009) and broad-tailed hummingbird (*Selasphorus platycercus*; Hogan and Stoddard, 2018).

In studies of wild birds, measures of body condition are based on size, including body mass, linear measures of skeleton or wing size, or mass-adjusted linear size (Bribiesca et al., 2019; Freeman and Jackson, 1990; Labocha and Hayes, 2012; Peig and Green, 2009; Peig and Green, 2010). Here, we show that body mass and body size variation within breeding male ruby-throated hummingbirds is not correlated with their peak flight muscle performance (Fig. 1B and Fig. S4). Compared to other bird families, hummingbirds undergo proportionately large fluctuations in body mass within individuals owing to their frequent nectar meals (Carpenter et al., 1983; Carpenter et al., 1991; Carpenter et al., 1993). In many other species, individuals may harbour substantial among-individual differences in muscle capacity that may be undetectable from standard measures of morphological size.

### Conclusion

Here, we show that peak performance on the load-lifting assay is highly repeatable in male ruby-throated hummingbirds, with large among-individual differences. Furthermore, our results indicate that these individual differences in burst muscle capacity are unrelated to body size variation in this species, and that peak performance may reveal important differences in body condition that are not captured by other methods. Our simulation results indicate that robust performance assays can accurately measure underlying individual maximal performance, even with a single measurement. Nevertheless, sampling individuals longitudinally can improve accuracy for studies that seek to measure individual differences in maximal ability.

## Acknowledgements

We would like to thank Nour Salama, Zoe Waller, and the Queen’s University Biological Station. We also thank Greg Bulté, Bruce McKay, Charlie and Sarah Anne Szabototh, and the Newlands and Bachhuber families.

## Data and resource availability

Data and code are available at: https://osf.io/38x9u/overview?view_only=153deeac777948f19df733707ab3ffbb.

