## supplement for "Peak performance is repeatable and captures large individual differences in ruby-throated hummingbirds"

**Movie 1. Load-lifting trial with ruby-throated hummingbirds.** The first two sequences show the positioning of the bird and loaded thread, and a series of lifts accomplished immediately after release in the cylinder. The third sequence shows how the experimenter’s presence and movement could also elicit a lift.

*Literature review of previous animal load-lifting studies*

On August 11^th^ 2026, we entered the following prompt in the Web of Science search engine: TS = ((((("Load" OR "mass" OR "weight") AND ("lift*")) OR ("max*" OR "peak" OR "burst") AND "power output") OR "burst muscle capacity") AND ("flight*" OR "fly*" OR "wing*") AND ("animal*" OR "species" OR "bird*" OR "bat*" OR "insect*") NOT ("human*" OR "human athlet*" OR "men" OR "man" or "women" OR "woman")). From this initial search, we narrowed the selection to include articles in English only. We screened through the 1,244 results, reading each title and abstract, and excluded papers that were irrelevant (e.g., free take-off flight performance assays). We read the selected papers (n = 43) and kept the ones that measured the maximal load-lifting capacity of a flying animal (n = 28). We added 5 other research articles from a separate Google Scholar search (n total = 33). Finally, we read each article, and supplemental material when present, to determine whether repeatability of load lifting was calculated. The literature review is summarized in Table S1 below.

Out of 33 studies where researchers measured the maximal load-lifting capacity of a flying animal, only two studies, both on other hummingbird species, reported examining multiple measures per individual (Table S1). Najar et al. (2026) repeated the load-lifting assay on black-chinned hummingbirds, *Archilochus alexandri*, with each bird tested repeatedly shortly after capture, and only a short rest (minutes) between repeated measures (Najar et al., 2026). Najar et al. estimated the repeatability of load-lifting performance to be 0.80 for males, and 0.75 when including males and females in this species (Najar et al., 2026). In a study of spatial memory in long-billed hermit hummingbirds, *Phaethornis longirostris*, Araya-Salas et al. (2018) conducted repeated load-lifting assays shortly after capture, and then re-captured and re-tested the same individual after an interval that ranged from hours to days or weeks apart. In their supplementary material, they estimate load-lifting repeatability ranging from 0.30 to 0.90, depending on the sample used in the analysis.

Thus, our finding that individual male ruby-throated hummingbirds differ extensively in peak load-lifting performance (repeatability 0.70) is consistent with the two previous studies of other hummingbird species. Furthermore, by systematically testing individuals within and across days under the same conditions, we establish here that an additional 23% of variation can be explained by short-term fluctuations in a bird’s performance.

*Designing the load-lifting apparatus*

The original load-lifting apparatus was a hanging cylinder (~10 cm above ground as per Chai and colleagues, 1997) with evenly perforated walls. Other studies have used solid-walled chambers (eg., Altshuler, 2006). During preliminary testing, we noticed that birds seemed distracted by the perforated wall design, not flying vertically, but rather from side to side. The final design with opaque walls and no hanging space at the bottom resulted in much more consistency in birds flying vertically towards the open top of the cylinder.

During the preliminary stages of the study design, we also tested on a small number of individuals whether letting the bird escape freely from the bottom of the cylinder (i.e., without a loaded thread attached) would improve its subsequent lifting performance. If experience of a successful escape led to an increase in load-lifting performance (as compared to performance before cylinder escape), it would suggest that the assay is not always eliciting maximal effort. We observed that an individual’s load-lifting performance before and after a cylinder escape remained the same, consistent with our expectation that the birds are consistently motivated to fly vertically with maximal effort in this assay.

*Scoring lifts*

To score the best lifts per trial, the experimenter reviewed each video twice: the first pass was done to identify the singular best lift (as ranked by the number of load units lifted), while the second pass was done to identify 5-10 lifts that were either equal or closely matched that individual best lift. We only scored lifts in which the thread was lifted vertically. Lifts in which the thread moved horizontally, caused by flailing, and lifts in which the thread touched the wall or the hand of the experimenter were not scored.

Because the number of initially scored lifts varied between trials, we retained the best five lifts as a consistent sample size per trial. When there were ties among the best lifts, we randomly selected among those tied best lifts. Using an example trial of RTHU_012 who lifted, in order, 29, 29, 29, 25, 27, 25 and 25 units: the five best in this trial included the three 29-unit lifts and the one 27-unit lift (giving us four best lifts), then we randomly selected one of the 25-unit lifts, to obtain his five best. This procedure of randomly selecting tied best lifts ensured that we sampled best lifts that occurred throughout the duration of the trial.

*Sources of variation in load-lifting performance*

In the ‘five best lifts’ protocol (*Lift_total_*), we report that birds tend to perform slightly better later within a trial (Table S4 and Fig. S3D). This trend, absent from the main ‘two best lifts’ protocol, was detectable in the ‘five best lifts’ protocol due to increased sample size of lifts per trial as compared to 'two best lifts’. Although it is a statistically significant effect, the effect size is very small (see Table S4 and Fig. S3D).

In the *Lift_relative_* analysis, we find that birds tended to achieve better *Lift_relative_* performance in later trials within a day, although the magnitude of this effect is again very small (Table S5). To investigate whether this association between *Lift_relative_* and trial-specific body mass could be driven by body mass loss over repeated trials within a day, we modelled per-trial body mass measurements with trial number as the fixed effect and bird ID as a random effect (Table S6). This analysis revealed that the body mass of the birds on test days decreased subtly across trials within a day (Table S6), which can explain why body-mass adjusted *Lift_relative_* performance increased subtly with trial number (Table S5), whereas we did not detect a change in a bird’s *Lift_total_* with trial number (Table 1 of the main text).

To examine whether skeletal body size predicted load lifting performance, we first used a principal component analysis (PCA) on three skeletal size measures (bill, keel, and tarsus lengths) that are positively correlated (Tables S2 and S3). We took the first component PC1, which accounted for 75% of the total variation, as a proxy of hummingbird skeletal body size (Table S3). We then performed a linear regression with the average individual *Lift_total_* performance as the response variable, and PC1 skeletal body size as the predictor (*n* = 13 values, one per male). We detected no effect of skeletal size on lifting performance in this analysis (coefficient estimate = -0.140 [95% CI -0.377, 0.097], *P*-value = 0.214; Fig. S4).

**Table S1. Literature review of previous animal load-lifting studies.** Results of animal load-lifting studies found on the Web of Science search engine, followed by a thorough screening. We screened through the results, reading the title, abstract, main text and supplementary material of each article, to identify which studies measured the maximal load-lifting capacity of a flying animal. From this final article selection, we noted the study species, whether the load-lifting assay was repeated, and whether the authors calculated the repeatability or not. Studies in which researchers measured flying animals’ maximal load-lifting capacity were retained, and information on repeatability calculation of the assay was extracted. Articles that were added from a separate Google Scholar search are marked with an asterisk (*).

| **Article** | **Study animal(s)** | **Repeated measures per individual** | **Repeatability estimate** |
| --- | --- | --- | --- |
| ***Altshuler, D. L.** (2006). Flight Performance and Competitive Displacement of Hummingbirds across Elevational Gradients. *The American Naturalist* **167**, 216–229. | 45 hummingbird species (Trochilidae family; *n* = 1-14 individuals per species) | No | - |
| **Altshuler, D. L. and Dudley, R.** (2003). Kinematics of hovering hummingbird flight along simulated and natural elevational gradients. *J Exp Biol* **206**, 3139–3147. | 45 hummingbird species (Trochilidae family; *n* = 1-14 individuals per species) | No | - |
| ***Altshuler, D. L., Stiles, F. G. and Dudley, R.** (2004a). Of Hummingbirds and Helicopters: Hovering Costs, Competitive Ability, and Foraging Strategies. *The American Naturalist* **163**, 16–25. | 3 hummingbird species: Fork-tailed nymph (*Thalurania furcata*; *n* = 9 males and 9 females), rufous hummingbird (*Selasphorus rufus*; *n* = 10 males and 14 females), and Koepcke’s hermit (*Phaethornis koepckeae*; *n* = 7 females and 10 males) | No | - |
| **Altshuler, D. L., Dudley, R. and McGuire, J. A.** (2004b). Resolution of a paradox: Hummingbird flight at high elevation does not come without a cost. *Proceedings of the National Academy of Sciences* **101**, 17731–17736. | 43 hummingbird species (Trochilidae family; *n* = 1-14 individuals per species) | No | - |
| **Altshuler, D. L., Welch, K. C., Jr, Cho, B. H., Welch, D. B., Lin, A. F., Dickson, W. B. and Dickinson, M. H.** (2010). Neuromuscular control of wingbeat kinematics in Anna’s hummingbirds (Calypte anna). *J Exp Biol* **213**, 2507–2514. | Anna’s hummingbird (*Calypte anna*; *n* = 4) | No | - |
| **Araya-Salas, M., Gonzalez-Gomez, P., Wojczulanis-Jakubas, K., López, V. and Wright, T. F.** (2018). Spatial memory is as important as weapon and body size for territorial ownership in a lekking hummingbird. *Sci Rep* **8**, 2001. | Long-billed hermits (*Phaethornis longirostris*; *n* = 7-24, depending on sub-sampling protocol) | **Yes**; repeated trials based on recapture opportunities (time between trial varies from a few hours to 36 days)^[1]^ | 0.30 - 0.90, depending on sub-sampling protocol |
| **Buchwald, R. and Dudley, R.** (2010). Limits to vertical force and power production in bumblebees (Hymenoptera: Bombus impatiens). *J Exp Biol* **213**, 426–432. | Bumblebee (*Bombus impatiens*; *n* = 62) | No | - |
| **Burnett, N. P., Keliher, E. L. and Combes, S. A.** (2022). An evaluation of common methods for comparing the scaling of vertical force production in flying insects. *Current Research in Insect Science* **2**, 100042. | 2 bee species: bumblebee (*Bombus impatiens*; *n* = 28), and orchard mason bee (*Osmia lignaria*; *n* = 25) | No | - |
| **Cano-Franco, A., Mancilla-Morales, M. D., Contreras-Rodríguez, A., Flores-Martínez, J. J., Gomez-Lunar, Z. and Ruiz, E. A.** (2025). Size and Sex Effects on Storm-Petrels’ Maximum Load-Lift at Takeoff. *Diversity* **17**, 417. | 2 bird species: least storm-petrel (*Hydrobates microsoma*, *n* = 48), and black storm-petrel (*Hydrobates melania*; *n* = 23) | No | - |
| **Chai, P. and Millard, D.** (1997). Flight and size constraints: hovering performance of large hummingbirds under maximal loading. *Journal of Experimental Biology* **200**, 2757–2763. | 4 hummingbird species: blue-throated hummingbird (*Lampornis clemenciae*; *n* = 2), Magnificent hummingbird (*Eugenes fulgens*; *n* = 3, black-chinned hummingbird (*Archilochus alexandri*; *n* = 5), and rufous hummingbird (*Selaphorus rufusl*; *n* = 1) | No | - |
| **Chai, P., Chen, J. S. C. and Dudley, R.** (1997). Transient Hovering Performance of Hummingbirds Under Conditions of Maximal Loading. *Journal of Experimental Biology* **200**, 921–929. | Ruby-throated hummingbird (*Archilochus colubris*; *n* = 6) | No | - |
| ***Dakin, R., Segre, P. S., Straw, A. D. and Altshuler, D. L.** (2018). Morphology, muscle capacity, skill, and maneuvering ability in hummingbirds. *Science* **359**, 653–657. | 25 hummingbird species (263 individuals total across species) | No | - |
| **Davidovich, H. and Ribak, G.** (2018). Loaded flight in male *Ischnura elegans* and its relationship to copulatory flight. *Journal of Insect Physiology* **110**, 44–56. | Blue-tailed damselfly (*Ischnura elegans*; *n* = 19) | No | - |
| ***Davis, R. and Cockrum, E. L.** (1965). Experimentally Determined Weight Lifting Capacity in Individuals of Five Species of Western Bats. *J Mammal* **45**, 643–644. | 5 bats species (*n* = 1-3 individuals per species) | No | - |
| **Dillon, M. E. and Dudley, R.** (2004). Allometry of maximum vertical force production during hovering flight of neotropical orchid bees (Apidae: Euglossini). *J Exp Biol* **207**, 417–425. | 8 orchid bee species (*n* = 3-29 individuals per species) | No | - |
| **Fernández, M. J., Driver, M. E. and Hedrick, T. L.** (2017). Asymmetry costs: effects of wing damage on hovering flight performance in the hawkmoth Manduca sexta. *J Exp Biol* **220**, 3649–3656. | Hawkmoth (*Manduca sexta*; *n* = 8) | No | - |
| **Groom, D. J. E., Toledo, M. C. B. and Welch, K. C.** (2017). Wingbeat kinematics and energetics during weightlifting in hovering hummingbirds across an elevational gradient. *J Comp Physiol B* **187**, 165–182. | 4 hummingbird species: (*Clytolaema rubricauda*; *n* = 13), (*Leucochloris albicollis*, *n* = 8), (*Thalurania glaucopis*, *n* = 15), and (*Lophornis chalybeus*, *n* = 4) | No | - |
| **Johnson, S. A. and Cartar, R. V.** (2014). Wing wear, but not asymmetry in wear, affects load-lifting capability in bumble bees Bombus impatiens. *Canadian Journal of Zoology* **92**, 179–184. | Bumble bee (*Bombus impatiens*; *n* = 74) | No | - |
| **Kou, G., Wang, Y., Dudley, R., Wu, Y. and Li, D.** (2022). Coping with captivity: takeoff speed and load-lifting capacity are unaffected by substantial changes in body condition for a passerine bird. *Journal of Experimental Biology* **225**,. | Eurasian tree sparrow (*Passer montanus*; *n* = 40) | No | - |
| **Kou, G., Wang, Y., Ge, S., Yin, Y., Sun, Y. and Li, D.** (2023). Moderate mass loss enhances flight performance via alteration of flight kinematics and postures in a passerine bird. *J Exp Biol* **226**, jeb245862. | Eurasian tree sparrow (*Passer montanus*; *n* = 17) | No | - |
| **Lehmann, F.-O.** (1999). Ambient temperature affects free-flight performance in the fruit fly Drosophila melanogaster. *J Comp Physiol B* **169**, 165–171. | Fruit fly (*Drosophila melanogaster*; *n* = 92) | No | - |
| **Mahalingam, S. and Welch, K. C., Jr** (2013). Neuromuscular control of hovering wingbeat kinematics in response to distinct flight challenges in the ruby-throated hummingbird, Archilochus colubris. *Journal of Experimental Biology* **216**, 4161–4171. | Ruby-throated hummingbird (*Archilochus colubris*; *n* = 4) | No | - |
| **Marden, J. H.** (1987). Maximum Lift Production During Takeoff in Flying Animals. *J Exp Biol* **130**, 235–258. | 49 insect species (1-12 individuals per species), 9 bird species (1-2 individuals per species) and 3 bat species (2-3 individuals per species) | No | - |
| **Mountcastle, A. M. and Combes, S. A.** (2013). Wing flexibility enhances load-lifting capacity in bumblebees. *Proc Biol Sci* **280**, 20130531. | Bumble bee (*Bombus impatiens*; *n* = 14) | No | - |
| ***Najar, N. A., Wilcox, S. C., Fernandez, L. R. and Clark, C. J.** (2026). Effects of wingbeat kinematics and muscle size on maximum flight speed in hummingbirds. *J Exp Biol* jeb.251012. | Black-chinned hummingbird (*Archilochus alexandri*; *n* = 53) | **Yes**; four trials in one day (one-minute rest period between trials), and kept the best 3 trials only | 0.73 - 0.80, depending on sex |
| **Ortega-Jimenez, V. M., Alvarez-Borrego, S., Arriaga-Ramirez, S., Bridge, E. S. and Renner, M.** (2011). Maximum Load-Carrying During Takeoff of Leach’s Storm-Petrel Oceanodroma leucorhoa and Cassin’s Auklet Ptychoramphus aleuticus. *cowa* **34**, 102–106. | Cassin’s auklet (*Ptychoramphus aleuticus*, *n* = 19), and leach’s storm-petrel (*Oceanodroma leucorhoa*; *n* = 26) | No | - |
| **Roberts, J. C. and Cartar, R. V.** (2015). Shape of wing wear fails to affect load lifting in common eastern bumble bees (Bombus impatiens) with experimental wing wear. *Can. J. Zool.* **93**, 531–537. | Bumblebee (*Bombus impatiens*; *n* = 57) | No | - |
| **Samejima, Y. and Tsubaki, Y.** (2010). Body temperature and body size affect flight performance in a damselfly. *Behav Ecol Sociobiol* **64**, 685–692. | Japanese damselfly(*Mnais costalis*; *n* = 30) | No | - |
| **Segre, P. S., Dakin, R. and Altshuler, D. L.** (2025). Turning trade-offs: hummingbird power reserves are used to decrease turning radius or increase turning velocity. *Journal of Zoology* **326**, 201–206. | Anna’s hummingbird (*Calypte anna*; *n* = 20) | No | - |
| **Segre, P. S., Dakin, R., Zordan, V. B., Dickinson, M. H., Straw, A. D. and Altshuler, D. L.** (2015). Burst muscle performance predicts the speed, acceleration, and turning performance of Anna’s hummingbirds. *eLife* **4**, e11159. | Anna’s hummingbird (*Calypte anna*; *n* = 20) | No | - |
| **Sun, Y.-F., Ren, Z.-P., Wu, Y.-F., Lei, F.-M., Dudley, R. and Li, D.-M.** (2016). Flying high: limits to flight performance by sparrows on the Qinghai-Tibet Plateau. *J Exp Biol* **219**, 3642–3648. | Eurasian tree sparrow (*Passer montanus*; *n* = 52) | No | - |
| **Wang, Y., Yin, Y., Ge, S., Li, M., Zhang, Q., Li, J., Wu, Y., Li, D. and Dudley, R.** (2019). Limits to load-lifting performance in a passerine bird: the effects of intraspecific variation in morphological and kinematic parameters. *PeerJ* **7**, e8048. | Eurasian tree sparrow (*Passer montanus*; *n* = 52) | No | - |
| **Wilcox, S. C. and Clark, C. J.** (2025). Flight performance during courtship in male black-chinned hummingbirds, *Archilochus alexandri*. *Animal Behaviour* **230**, 123368. | Black-chinned hummingbird (*Archilochus alexandri*; *n* = 18) | No | - |

**Table S2. Pearson’s correlation coefficients for three skeletal size measures.** Bill length, keel length, and tarsus length were all positively correlated (*n* = 13). Each value in the table reports the Pearson’s correlation result from the column and row measurements.

|  | **Bill length** | **Keel length** | **Tarsus length** |
| --- | --- | --- | --- |
| **Bill length** | - | 0.58 | 0.69 |
| **Keel length** |  | - | 0.58 |
| **Tarsus length** |  |  | - |

**Table S3. Principal components analysis used to quantify skeletal body size.** Each variable was standardized prior to running the analysis. The first row shows the cumulative proportion of variance explained by each component in percentage (Cum. Var. %), and the last three rows show the loading from each component for bill, keel and tarsus lengths.

|  | **Component 1** | **Component 2** | **Component 3** |
| --- | --- | --- | --- |
| **Cum. Var. %** | 75% | 90% | 100% |
| **Bill loading** | -0.59 | 0.40 | 0.70 |
| **Keel loading** | -0.55 | -0.83 | 0.01 |
| **Tarsus loading** | -0.59 | 0.38 | -0.71 |

**Table S4. Analysis of lifting performance using the five best lifts per trial.** The dependent variable is *Lift_total_*. All fixed effect predictors were centred and standardized, so that the coefficient estimates here are comparable. Variables are bolded when *P*-values are below 0.05.

| Fixed effect predictor | Estimate | 95% CIs [lower, upper] | *P*-value |
| --- | --- | --- | --- |
| Body mass_avg trial_ | 0.013 | [-0.092, 0.129] | 0.813 |
| Day number | 0.020 | [-0.059, 0.097] | 0.631 |
| Trial number | 0.033 | [-0.048, 0.114] | 0.427 |
| Time in day | -0.064 | [-0.175, 0.051] | 0.280 |
| **Time within trial** | **0.048** | **[0.023, 0.072]** | **<0.001** |
| Random effect | Proportion of variance (%) | 95% CIs [lower, upper] | *P*-value |
| **Bird ID** | **67** | **[38, 81]** | **<0.0001** |
| **Bird ID * day** | **9** | **[3, 22]** | **0.0002** |
| **Bird ID * day * trial** | **9** | **[5, 17]** | **<0.0001** |

**Table S5. Analysis of lifting performance using the two best lifts per trial of relative mass lifted.** The dependent variable is *Lift_relative_*. All fixed effect predictors were centred and standardized, so that the coefficient estimates here are comparable. Variables are bolded when *P*-values are below 0.05.

| Fixed effect predictor | Estimate | 95% CIs [lower, upper] | *P*-value |
| --- | --- | --- | --- |
| Day number | 0.014 | [-0.013, 0.041] | 0.314 |
| **Trial number** | **0.041** | **[0.015, 0.066]** | **0.003** |
| Time in day | -0.023 | [-0.061, 0.017] | 0.258 |
| Time within trial | 0.003 | [-0.008, 0.013] | 0.624 |
| Random effect | Proportion of variance (%) | 95% CIs [lower, upper] | *P*-value |
| **Bird ID** | **71** | **[45, 84]** | **<0.0001** |
| **Bird ID * day** | **7** | **[1, 21]** | **0.0042** |
| **Bird ID * day * trial** | **15** | **[8, 29]** | **<0.0001** |

**Table S6. Analysis of hummingbird body mass fluctuations across trials.** The dependent variable is the trial-average individual body mass in grams (average of before and after trial). Variables are bolded when *P*-values are below 0.05.

| Fixed effect predictor | Estimate | 95% CIs [lower, upper] | *P*-value |
| --- | --- | --- | --- |
| **Trial number** | **-0.047** | **[-0.055, -0.039]** | **<0.0001** |
| Random effect | Proportion of variance (%) | 95% CIs [lower, upper] | *P*-value |
| **Bird ID** | **77** | **[54, 87]** | **<0.0001** |

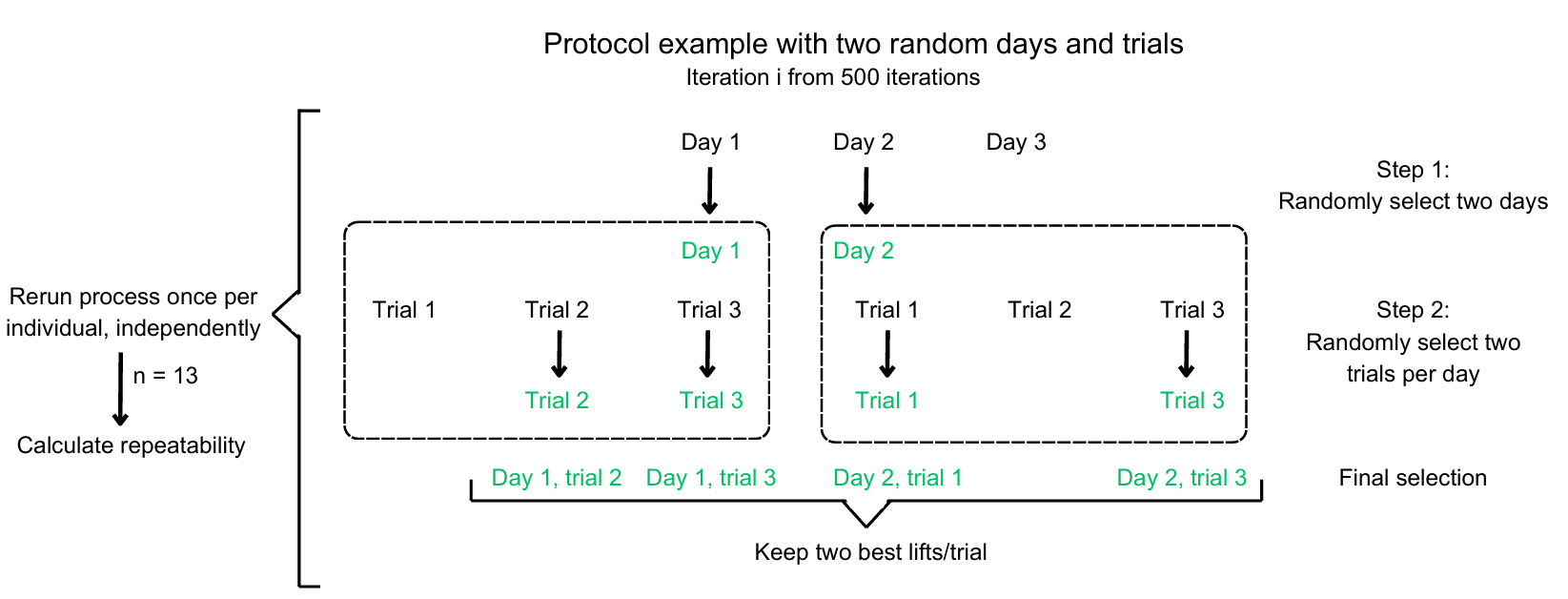

**Fig S1. Sub-sampling method used to generate different sampling protocol scenarios.** The example here shows how we generated a protocol with two test days and two trials per day. In step 1, two days are randomly sampled (of three), and in step 2, two trials are randomly selected from each day. The green text shows a specific result for one individual, and the process is repeated for each individual, independently.

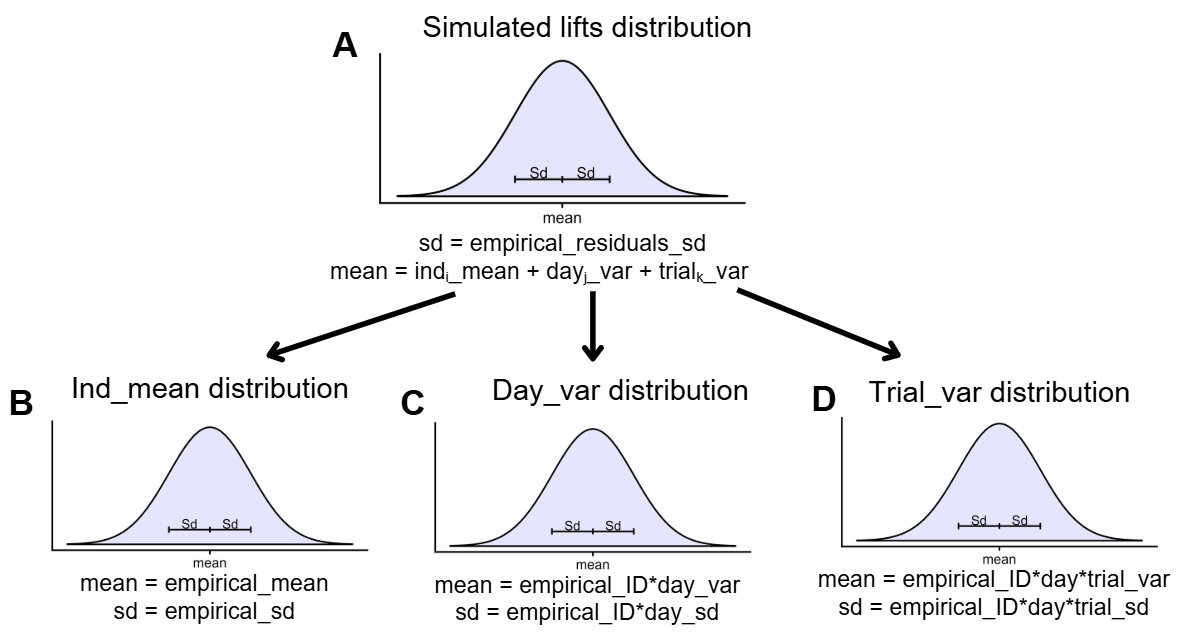

**Fig. S2. Simulation testing how sampling protocol affects the measurement of individual maxima**. Sources of variation in simulated lift performance are normally distributed, with their respective means and standard deviations as shown (*sd* on the figure). For each *in silico* individual (*n* = 13), a mean was randomly sampled from a normal distribution (B) with mean and standard deviation derived from the empirical hummingbird data. For each simulated day of trials (*n* = 3 days per bird, and *n* = 3 trials per bird), we added a daily fluctuation (C) and a trial-specific fluctuation (D) to that individual’s mean performance (A). Those daily and trial fluctuations were randomly sampled from normal distributions with their respective means and standard deviations derived from the empirical data. Simulated lifts were randomly sampled (*n* = 5 per trial) from the resulting normal distribution (A) for a given bird, day and trial, with residual error derived from the empirical data.

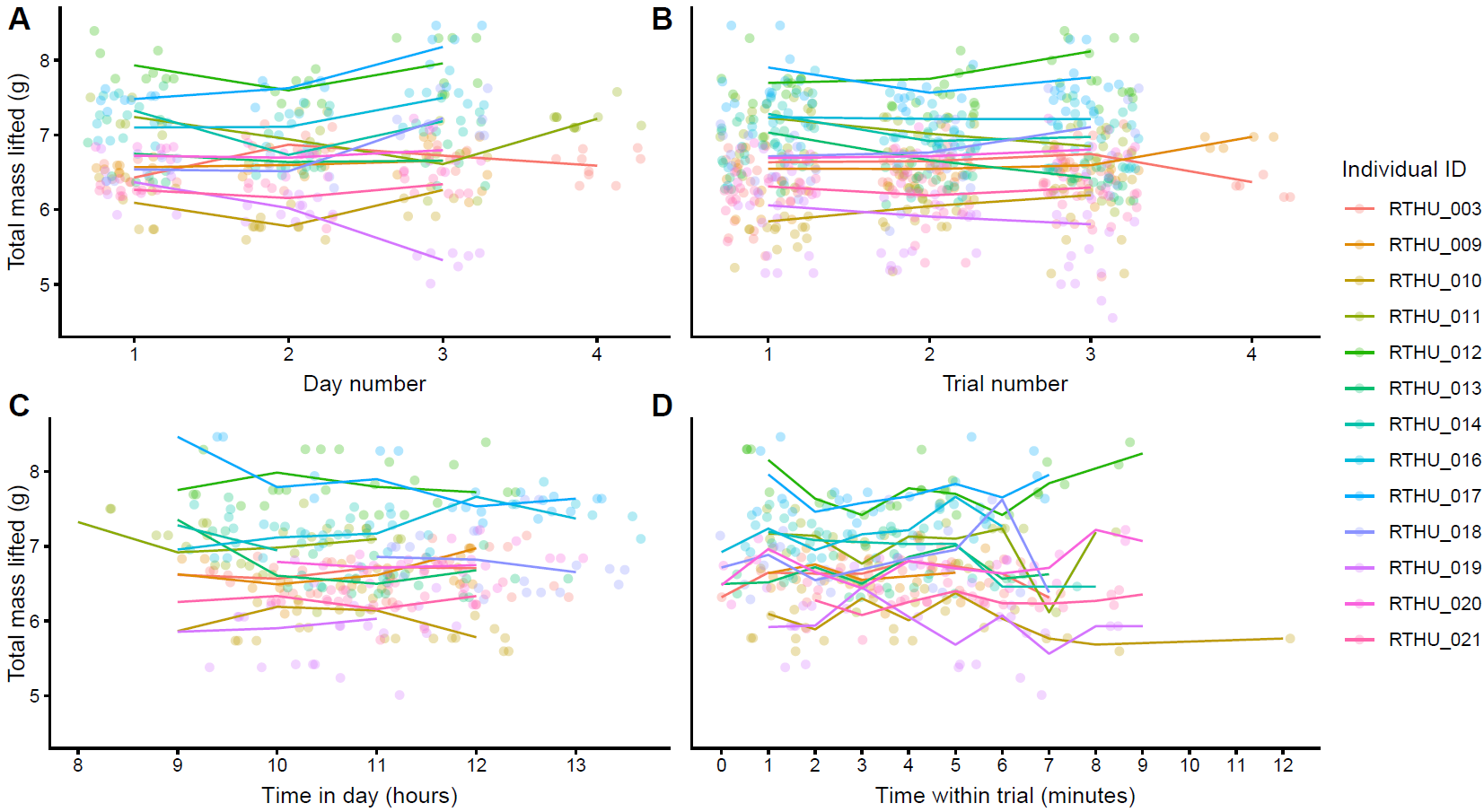

**Fig S3. Effects of experience and time on load-lifting performance.** Total lifted mass (*Lift_total_*) in relation to (A) day number, (B) trial number (within a day), (C) time in the day, and (D) time within a trial. Each point represents a recorded lift, colored by bird ID. For clarity, only the two highest-performance lifts per trial are shown here. The lines connect means from each male, which are rounded to the nearest hour in (C) and are rounded to the nearest minute in (D). In (A) and (B), points are jittered along the x axis for ease of visualizing the distributions.

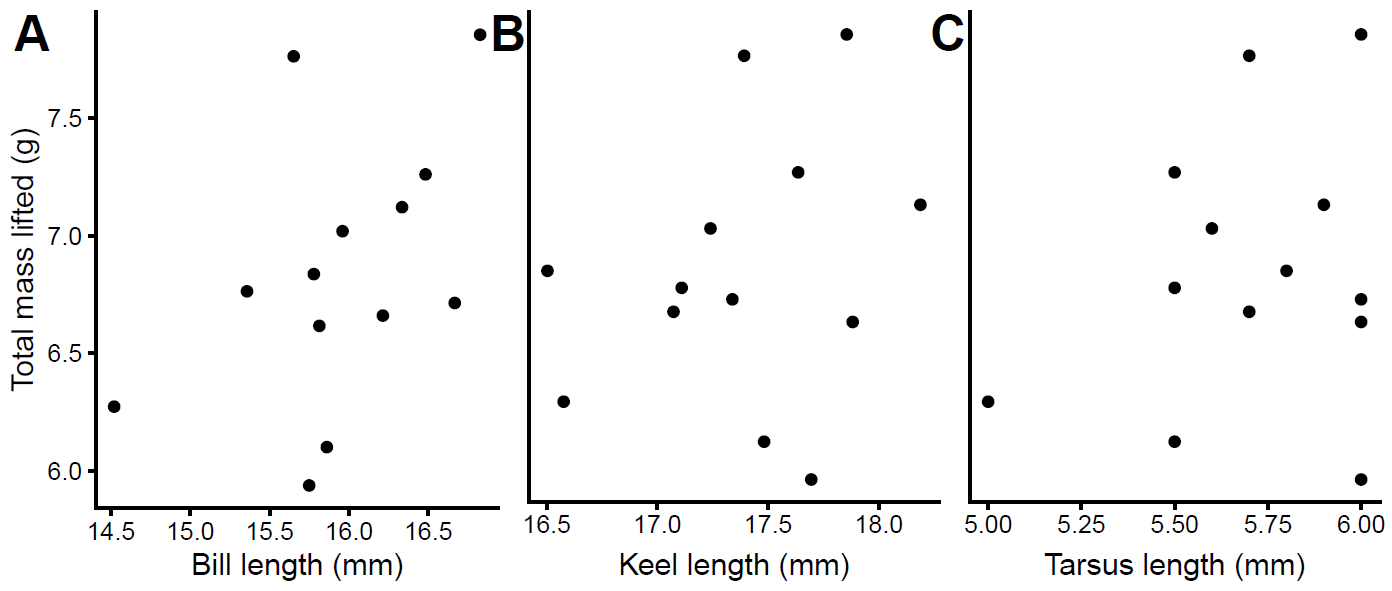

**Fig S4. Relationship between load-lifting performance and (skeletal) body size measurements.** Average loading-lifting performance (*Lift_total_*) of 13 hummingbird males plotted as a function of (A) bill length, (B) keel length and (C) tarsus length.

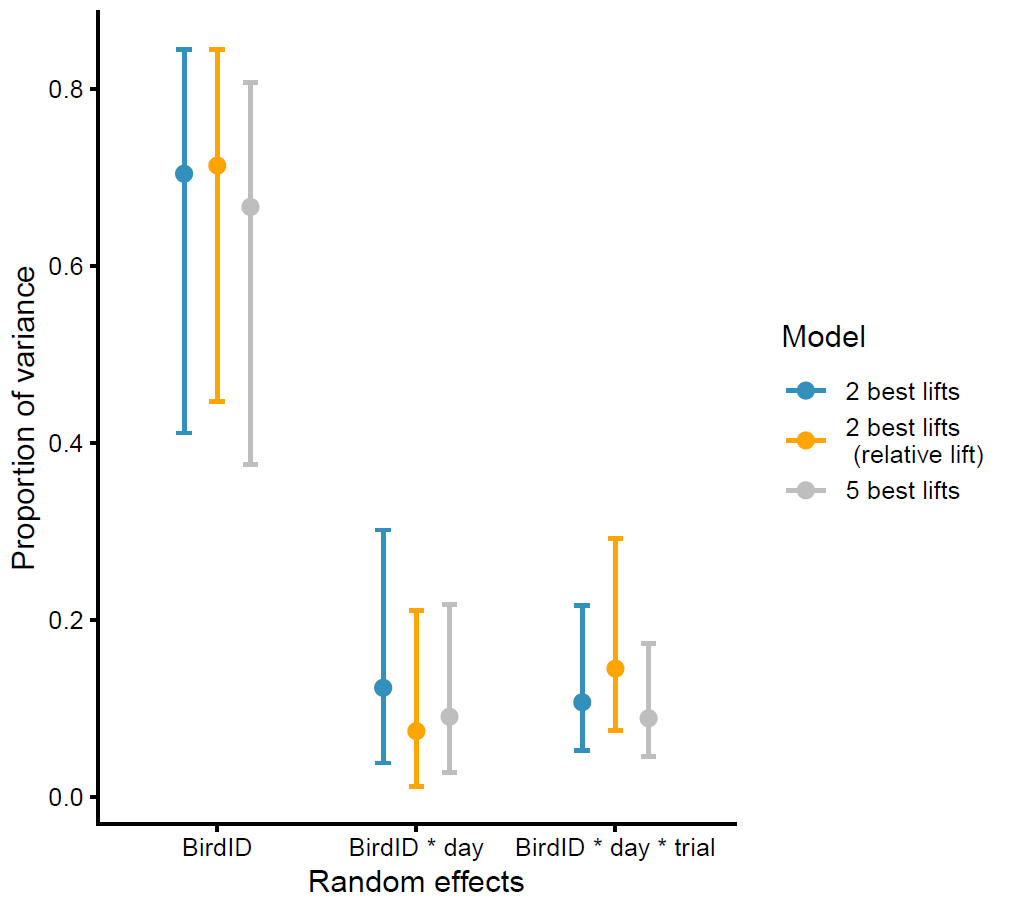

**Fig S5. Proportion of variance in peak lifting performance explained by individual differences (BirdID), day-to-day differences (BirdID * day), and trial-to-trial differences (BirdID * day * trial).** Estimates derived from different models are compared here. The “2 best lifts” model accounted for effects of bird body mass, day order, trial order, time of day, and time within a trial, using the two best total-mass lifts per trial (*Lift_total_*). The “2 best lifts (relative lifts)” model accounted effects of day order, trial order, time of day, and time within a trial, using the two best mass-relative lifts per trial (*Lift_relative_*). The “5 best lifts” model accounted for effects of bird body mass, day order, trial order, time of day, and time within a trial, using the five best total-mass lifts per trial (*Lift_total_*). Repeatability is the proportion of variance attributed to Bird ID. Error bars show 95% confidence intervals from bootstrapping (*n* = 1,000 iterations).

1. Sample sizes and time between trials were determined from the authors’ shared unpublished data
